# Chemotherapy causes a reversible decrease in *VMP1/MIR21* DNA methylation in granulocytes from breast cancer survivors

**DOI:** 10.1101/2024.08.15.608076

**Authors:** Marie-Louise Abrahamsen, Zuzanna Jachowicz, Kristian Buch-Larsen, Djordje Marina, Michael Andersson, Peter Schwarz, Flemming Dela, Linn Gillberg

**Author notes:** **Correspondence** Linn Gillberg, Department of Biomedical Sciences, University of Copenhagen, Blegdamsvej 3B, 2200 Copenhagen, Denmark. **Author contribution** M-L.A. and L.G. designed the research. K.B. and D.M. recruited the patients. P.S., F.D., and L.G. provided resources. M-L.A., Z.J., and L.G. performed data acquisition, analysis, and interpretation. M-L.A. and L.G. wrote the manuscript. All authors critically revised the manuscript and approved the final version of the paper to be published. L.G. takes responsibility for all aspects of the reliability of the presented data. **Ethical approval, patient consent and clinical trial registration** The study was conducted in accordance with the Declaration of Helsinki and approved by the Ethics Committee of The Capital Region, Denmark (project number H-18016600 with amendments 67762 and 74303) and the Data Protection Agency. Informed consent was obtained from all subjects enrolled in the study. The clinical trial (registration number NCT03784651) was registered on www.clinicaltrials.gov on December 24, 2018.

## Abstract

**Background:** DNA methylation alterations within the *VMP1/MIR21* gene region, a potential epigenetic biomarker of systemic inflammation, have been demonstrated in mononuclear blood cells from early breast cancer (BC) patients after chemotherapy. Whether these changes are present in granulocytes, persist in the years after treatment, and affect *VMP1* or *MIR21* gene expression, remains unknown.

**Aim:** We aimed to investigate whether adjuvant chemotherapy alters the DNA methylation and gene expression of *VMP1*/*MIR21* in granulocytes from postmenopausal BC patients and, if so, whether these treatment-induced changes are reversible in the first two years after completed chemotherapy.

**Methods:** Whole blood samples were obtained from 30 postmenopausal BC patients before chemotherapy and every six months for two years, and from 10 healthy age- and BMI-matched controls. DNA and RNA was extracted from isolated granulocytes, and DNA methylation of four CpG sites located in the gene body of *VMP1*, which is situated in the promoter region of *MIR21*, was assessed through bisulfite pyrosequencing. qPCR was used for assessment of *VMP1* and *MIR21* expression.

**Results:** *VMP1/MIR21* was significantly demethylated in granulocytes from BC patients shortly after completed chemotherapy compared to before (10 percentage points decrease, *p*<0.0001). Six months thereafter, DNA methylation values were significantly increased (6 percentage points, p = 0.002), and they were further increased to pre-chemotherapy levels 12, 18 and 24 months post chemotherapy. Chemotherapy did not cause significant changes in the expression of *VMP1* or *MIR21*.

**Conclusion:** The unique follow-up samples in this study demonstrated that chemotherapy induced a transient reduction in DNA methylation of the *VMP1/MIR21* region in granulocytes from postmenopausal BC patients. Although transient, chemotherapy-induced epigenetic changes in blood cells may contribute to the increased risk of inflammatory-related comorbidities in BC survivors.

## 1. Introduction

Breast cancer (BC) is the most common cancer worldwide, however, earlier diagnosis and improved treatments have resulted in an increasing group of BC survivors. The treatment for BC is often multimodal consisting of surgery followed by adjuvant treatments, which may include chemotherapy, radiotherapy, and anti-estrogen treatment to maximize the therapeutic efficacy and minimize the risk of cancer recurrence. However, previous studies have reported long-term side effects after the extensive adjuvant treatments, including chronic inflammation^1–3^. This increases the risk of inflammatory-related comorbidities in BC survivors, but only limited data are accessible on the molecular mechanisms behind these side effects.

It has been demonstrated that chemotherapy can lead to DNA methylation alterations in healthy cells of BC patients, including isolated blood cells and whole blood^4–7^. DNA methylation is the most studied epigenetic modification and refers to the methylation of the fifth position of the cytosine base to produce 5-methylcytosine. Methylation of cytosines is largely restricted to cytosines immediately upstream from guanine, and these sites are known as CpG sites. DNA methylation of CpG sites in e.g. the promoter region can interfere with transcription of the gene or micro RNAs (miRNA).

Yao et al. demonstrated that chemotherapy altered the DNA methylation landscape of blood leukocytes from BC patients. The most significant change was the demethylation of five CpG sites in exon 11 of *VMP1*, encoding Vacuole Membrane Protein 1^4^. Chemotherapy-induced demethylation of *VMP1* was also demonstrated by Smith et al. who investigated DNA methylation in peripheral blood mononuclear cells (PBMCs) from chemotherapy versus non-chemotherapy-treated BC patients. Furthermore, this study showed that demethylation of *VMP1* was associated with increased plasma levels of pro-inflammatory cytokines^5^. This is in line with results from previous epigenome-wide association studies, where demethylation of *VMP1* was shown to correlate with inflammation-related phenotypes and conditions^8–13^, indicating that the *VMP1* locus could be a potential epigenetic biomarker of inflammation. Moreover, the *VMP1* locus lies within the promoter region of *MIR21*^14^ encoding miR-21, and this miRNA has been shown to be involved in pro-inflammatory responses^15^. Importantly, the decreased DNA methylation of *VMP1/MIR21* has been shown to be associated with an increase in miR-21 expression in whole blood^14^ and PBMCs^5^, demonstrating a link between the gene expression and DNA methylation at this locus. Furthermore, the expression of miR-21 has been shown to negatively correlate with *VMP1* expression in colorectal cancer tissues^16^, establishing a relation between miR-21 and *VMP1* expression.

Although two independent studies have demonstrated that chemotherapy alters the DNA methylation of *VMP1/MIR21* in blood cells from BC patients, none of the studies have a follow-up time longer than six months post chemotherapy. Here, we aimed to investigate if *VMP1/MIR21* DNA methylation changes with chemotherapy in granulocytes from postmenopausal early BC patients and whether potential changes are reversible or persist in the first two years post chemotherapy. In addition, we aimed to investigate whether adjuvant chemotherapy alters the gene expression of *VMP1* and *MIR21*, and whether transcripts correlate with the DNA methylation of the *VMP1/MIR21* region.

## 2. Methods and materials

### 2.1 Subjects and blood sampling

Thirty postmenopausal BC patients aged 50-70 years eligible for adjuvant chemotherapy were included in this study. They were all diagnosed with early BC (stage I-III) and recruited to the clinical study *Healthy Living after Breast Cancer* (NCT03784651) at the Department of Oncology, Rigshospitalet, Copenhagen, Denmark^17^. Exclusion criteria were preexisting endocrine or metabolic diseases and malignancy prior to the current BC diagnosis. Furthermore, 10 age- and BMI-matched controls were included in the study (**Supplementary table 1**). Blood sampling of BC patients was performed at Rigshospitalet before initiation of adjuvant chemotherapy and every six months in the first two years after completed chemotherapy (i.e. at their 6-, 12-, 18-, and 24-months visits). Seven patients had completed their 24 months visit when examinations described below were initiated. Control participants were examined at baseline. The patients’ tumor characteristics, surgery, and antineoplastic treatment regimens are shown in **Supplementary table 2**.

### 2.2 Isolation of granulocytes from whole blood

Venous whole blood (9 or 15 mL) was sampled in K_2_EDTA tubes and diluted to a final volume of 30 mL with phosphate buffered saline (PBS; Gibco™, Thermo Fisher Scientific, Waltham, MS, USA). The diluted blood was carefully layered on top of 15 mL Lymphoprep™ density gradient medium (StemCell Technologies Inc., Vancouver, Canada), and centrifuged at 800xg for 15 minutes (room temperature, acceleration 9, deceleration 2) to separate granulocytes and erythrocytes from PBMCs. After collection of the PBMCs, the granulocyte and erythrocyte layer was collected with a Pasteur pipette and Hoffmann’s red blood cell lysis buffer (Region Hovedstadens Apotek, Copenhagen, Denmark) was added up to a final volume of 45 mL. Following 7 minutes of incubation at RT, the sample was centrifuged at 433xg for 5 minutes (room temperature, acceleration 9, deceleration 9) and the supernatant was discarded. The pellet was resuspended in 45 mL Hoffmann’s red blood cell lysis buffer and the incubation and centrifugation steps were repeated. After the second centrifugation, the granulocyte pellet was resuspended in 3 mL PBS and equally transferred to two 1.5 mL tubes followed by centrifugation for 3 minutes at 12,000xg. The supernatants were discarded, and the isolated granulocytes were kept at -80°C.

### 2.3 DNA and RNA isolation from granulocytes

DNA and RNA was extracted from isolated granulocytes using the magnetic bead-based KingFisher™ Duo Prime purification system (Thermo Fisher Scientific Inc., Massachusetts, USA), in combination with the Mag-Bind^®^ Total RNA 96 Kit and Mag-Bind^®^ Blood and Tissue DNA HDQ 96 Kit (Omega Bio-Tek Inc., Georgia, USA) following the manufactures guidelines. The final DNA and RNA concentrations were measured with the Qubit™ 4.0 Fluorometer (Thermo Fisher Scientific) using the Qubit dsDNA Broad Range Assay Kit and the Qubit RNA High Sensitivity Assay Kit (Invitrogen by Thermo Fisher Scientific, Waltham, MA, USA).

### 2.4 Bisulfite pyrosequencing of isolated granulocyte DNA

Bisulfite pyrosequencing is the golden standard for DNA methylation analysis of selected CpG sites and was used for investigation of DNA methylation levels. Bisulfide conversion of 300 ng genomic DNA was carried out using the EZ DNA Methylation-Lightning™ Kit (Zymo Research, California, USA). The pyrosequencing assay was designed in the PyroMark Assay Design 2.0 software (Qiagen, Hilden, Germany). A target region containing four specific CpG sites within a region of 108 base pairs in the gene body (exon 11) of *VMP1*, which are situated in the promoter region of *MIR21*, were covered (**Supplementary figure 1**). Primers were ordered from Tag Copenhagen (Frederiksberg, Denmark) with biotin bound to the 5’-end of the reverse PCR primer (Forward: 5’-TGGGTTATT GTATTTTGTTTTTAGTGT-3’, Reverse: 5’-Biotin-ACACTAACAAACCAACTTCACTTAT-3’, Sequencing: 5’-GTATTTTGTTTTTAGTGTTGTT-3’). Amplification of the bisulfite-converted DNA was carried out using the PyroMark PCR kit (Qiagen) and the designed primers. Prior to initiation of the pyrosequencing experiments, optimal conditions for the designed assay were defined. Pyrosequencing was performed with the PyroMark Q24 system, including PyroMark Q24 buffers, reagents, and vacuum workstation (Qiagen). The pyrosequencing results were evaluated with predefined criteria using the PyroMark Q24 software (Qiagen). After quality control, data was included for pre-chemo: n = 28, post-chemo: n = 28, 6 months: n = 21, 12 months: n = 16, 18 months: n = 8, 24 months: n = 7 and controls: n = 10. Average methylation levels of the four CpG sites analyzed are reported in this manuscript.

### 2.5 qPCR analysis

The isolated granulocyte RNA was used for gene expression analyses of *VMP1* and *MIR21*.

The *VMP1* gene expression was analyzed using the QuantiNova™ SYBR Green PCR kit with *ACTB* as an endogenous control, which has been validated as a reference gene for qPCR studies in human neutrophils^18^. The qPCR assays for *VMP1* (Forward: 5’-GACCAGAGACG TGTAGCAATG-3’, Reverse: 5’-ACAATGCTTTGACGATGCCATA-3’) and *ACTB* (Forward: 5’-CTGGAACGGT GAAGGTGACA-3’, Reverse: 5’-AAGGGACTTCCTGTAACAATGCA-3’) were ordered from Sigma Aldrich (Missouri, USA). cDNA synthesis was carried out using the QuantiTect Reverse Transcriptase Kit (Qiagen) following the manufactures guidelines.

The *MIR21* gene expression was investigated using the TaqMan^®^ Advanced miRNA Assay system (Applied Biosystems, Thermo Fisher Scientific, Waltham, Massachusetts, USA) with *MIR24* as an endogenous control, which has been shown to have a stable expression across most human tissues^19^. Primer assays (477992_mir for hsa-miR-24-3p and 477975_mir for hsa-miR-21-5p) were ordered from Applied Biosystems (Thermo Fisher Scientific). cDNA synthesis was carried out using the TaqMan Advanced miRNA cDNA synthesis kit (Thermo Fisher Scientific).

All samples were loaded in triplicates and negative controls were included in all runs. The qPCR was run on a QuantStudio 1 real-time cycler (QuantStudio™, Thermo Fisher). Using the ΔΔCt-method^20^, the fold difference in gene expression was calculated for the patients before chemotherapy relative to the controls, and the fold change for patients after chemotherapy was calculated relative to before chemotherapy. We only analyzed gene and miRNA expression in patient samples up to 12 months after chemotherapy.

### 2.6 Statistical analyses

The DNA methylation data was analyzed with unpaired t-tests (patients vs. controls) or with mixed-effects analysis with the Geisser-Greenhouse correction for variance and Tukey’s multiple comparisons test (patients at different visits). The gene expression data was analyzed using one-sample Wilcoxon test, where the median of the sample was compared with a hypothetical median set to be 1.0. Correlations between DNA methylation and gene expression were carried out with Spearman correlation using data from patients before treatment and controls. A difference was defined as significant if the p-value was p < 0.05, while a p-value of 0.05 ≤ p ≤ 0.10 defined a tendency.

## 3. Results

### 3.1 Adjuvant chemotherapy causes a reversible decrease in VMP1/MIR21 DNA methylation

The average DNA methylation of the *VMP1/MIR21* region decreased by 10 percentage points after chemotherapy compared to before (58% pre vs 48% post, **Figure 1A**, *p* < 0.0001). DNA methylation levels decreased in 23 out of the 26 BC patients that had measurements both before and after chemotherapy (**Figure 1B**). Interestingly, six months after completed chemotherapy the *VMP1/MIR21* methylation level had increased by 6 percentage points compared to shortly after treatment (53%, **Figure 1A**, *p* = 0.002). The DNA methylation level was further increased 12 months after chemotherapy compared to shortly after treatment (56%, **Figure 1A**, *p* = 0.0002), reaching a level that was no longer statistically different from before chemotherapy initiation (**Figure 1A**, *p* = 0.29). The DNA methylation levels 18 and 24 months after chemotherapy remained stable and similar to the level before chemotherapy initiation (59% and 56%, **Figure 1**).

**Figure 1:**
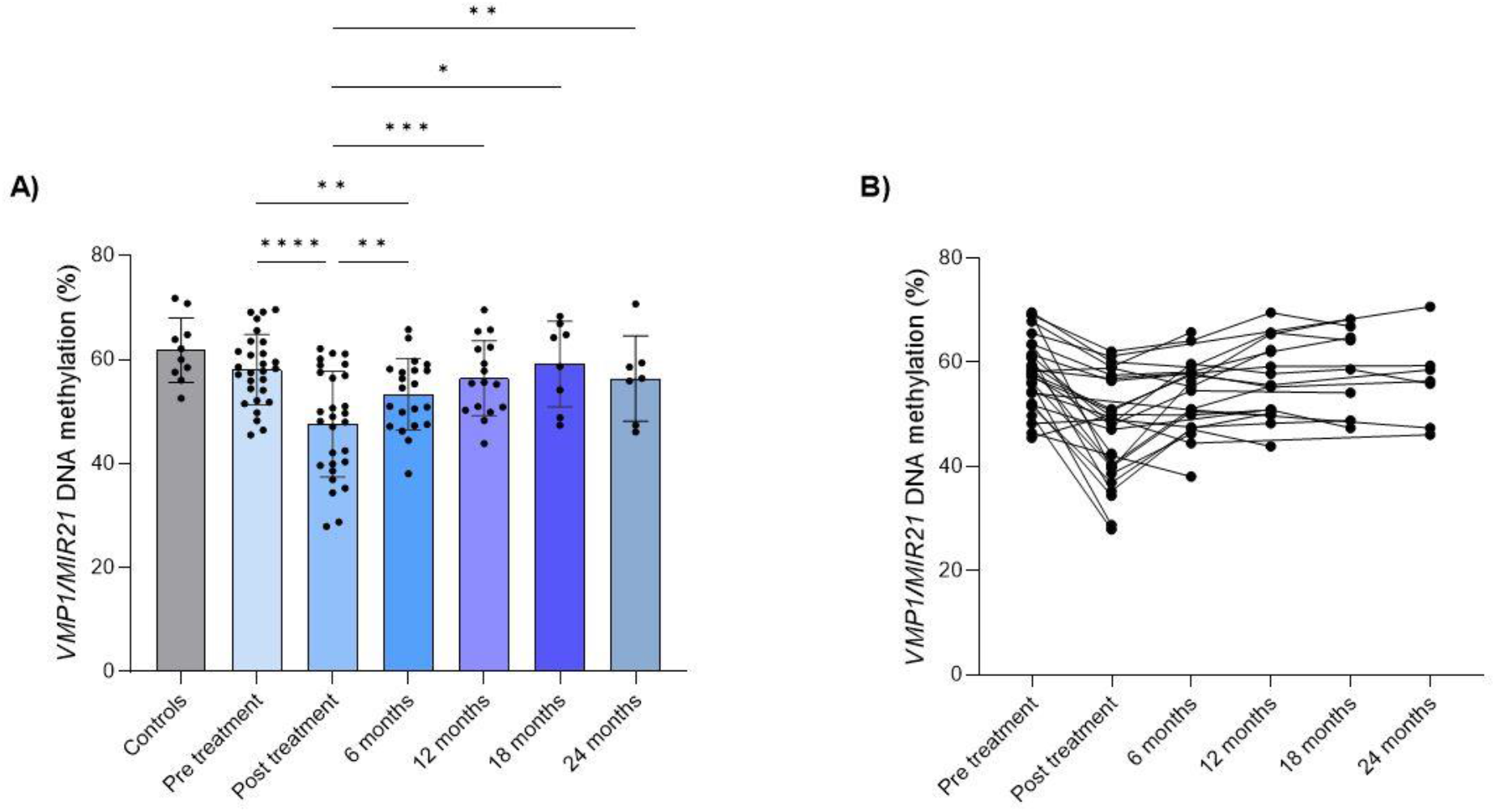
DNA Methylation of Four CpG Sites in *VMP1/MIR21* in Granulocytes from Healthy Controls and BC Patients. **A)** Average DNA methylation percentage of the four CpG sites analyzed in the gene body of *VMP1* and promoter region of *MIR21* in granulocytes from healthy controls and BC patients before chemotherapy, shortly after, 6 months, 12 months, 18 months, and 24 months after chemotherapy. Error bars show the standard deviation, whereas the top of the bars represents the mean. Dots represent each patient (pre: n = 28, post: n = 28, 6 months: n = 21, 12 months: n = 16, 18 months: n = 8, 24 months: n = 7) or control (n = 10). P-values are based on mixed-effects analysis with multiple comparisons. ****: p-value < 0.0001, ***: p-value < 0.001, **: p-value < 0.005, *: p-value < 0.05. **B)** Average DNA methylation percentage of the four CpG sites analyzed in *VMP1/MIR21* in granulocytes for each patient from before chemotherapy up to 24 months after chemotherapy. Dots represent each patient with lines connecting the same patient between visits.

Although not statistically significant (*p* = 0.13), the BC patients had on average 4% lower methylation levels than the controls before initiation of chemotherapy (**Figure 1A**).

### 3.2 The expression level of VMP1 and MIR21 did not change with chemotherapy

Since DNA methylation of the *VMP1/MIR21* locus has been shown to correlate with miR-21 levels^5,14^, and miR-21 levels has been shown to correlate with *VMP1* gene expression^16^, we analyzed *VMP1* and *MIR21* expression in the patient and control samples. In contrast to *VMP1/MIR21* DNA methylation levels, chemotherapy did not cause a significant change in *VMP1* or *MIR21* expression levels in granulocytes from the BC patients (**Figure 2B**, *VMP1 p* = 0.58, *MIR21 p* = 0.08). For *MIR21*, there was however a tendency in the patient group to a lower *MIR21* expression 6 and 12 months post chemotherapy compared to before (**Figure 3B**, *p* = 0.06 and *p* = 0.06). Furthermore, we found no difference in neither *VMP1* nor *MIR21* transcripts in granulocytes from BC patients before chemotherapy compared to healthy controls (**Figure 2A**, *VMP1 p* = 0.94, *MIR21 p* = 0.22). Finally, we found no correlation between average DNA methylation levels and *VMP1* gene expression levels (r = 0.02, *p* = 0.91) or *MIR21* levels (*MIR21* r = -0.26, *p* = 0.23). *MIR21* and *VMP1* expression levels did not intercorrelate (r = 0.24, *p* = 0.21).

**Figure 2:**
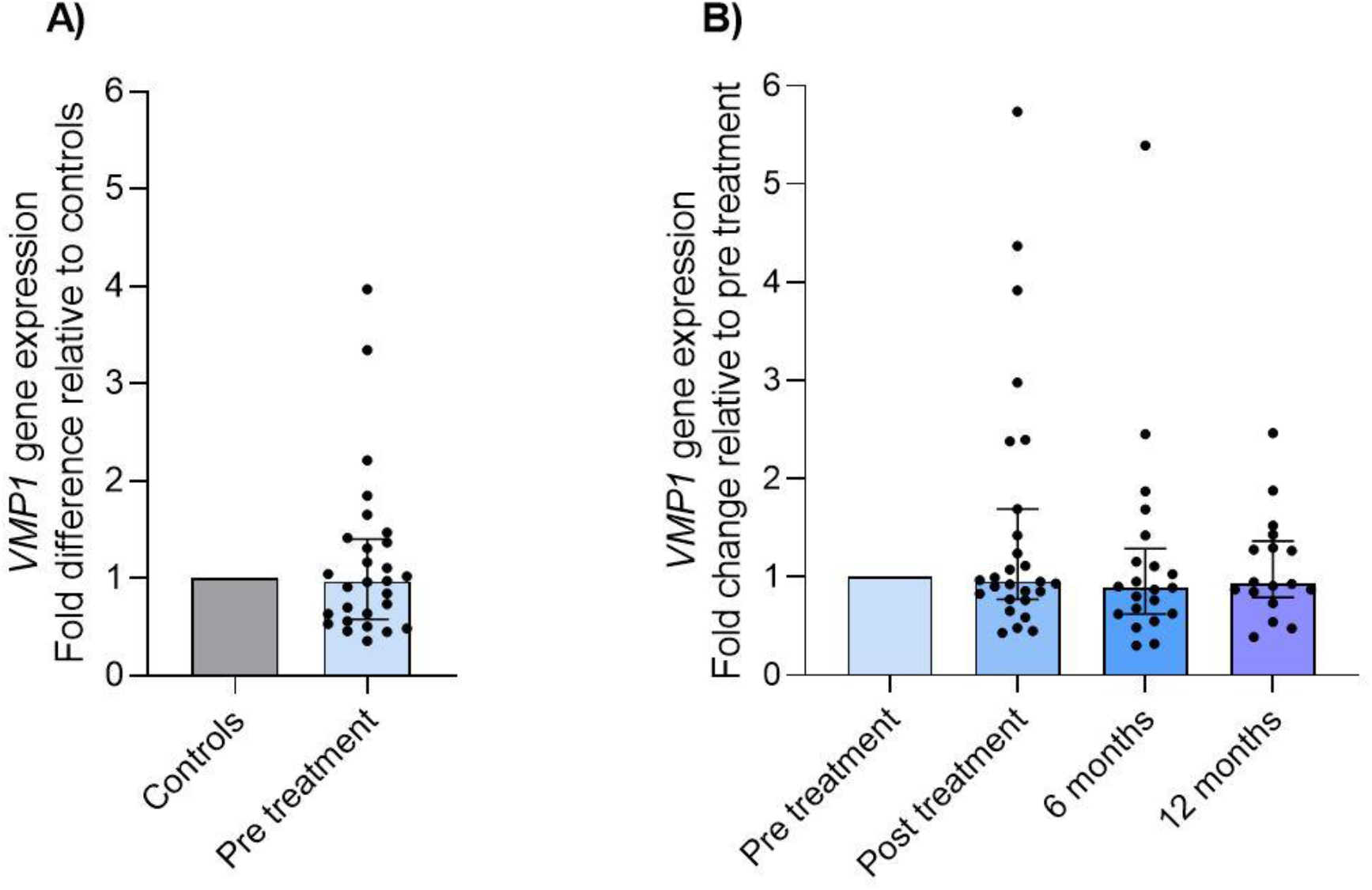
The Fold Difference or Change in Gene Expression of *VMP1* in Granulocytes from BC Patients and Healthy Controls. **A)** Fold difference in gene expression of *VMP1* in granulocytes from patients before chemotherapy normalized to the housekeeping gene *ACTB* and controls through calculation of ΔΔCt. Error bars show the interquartile range, whereas the top of the bars represents the medians. Controls are set to 1.0 as reference. Dots represent each patient (n = 28). **B)** Fold change in gene expression of *VMP1* in granulocytes from patients post chemotherapy, 6 months after chemotherapy, and 12 months after chemotherapy normalized to the housekeeping gene *ACTB* and patients before chemotherapy initiation (pre treatment) through calculation of ΔΔCt. Error bars show the interquartile range, whereas the top of the bars represents the medians. Patient data on pre treatment are set to 1.0 as reference. Dots represent each patient (post: n = 27, 6 months: n = 21, 12 months: n = 17).

**Figure 3:**
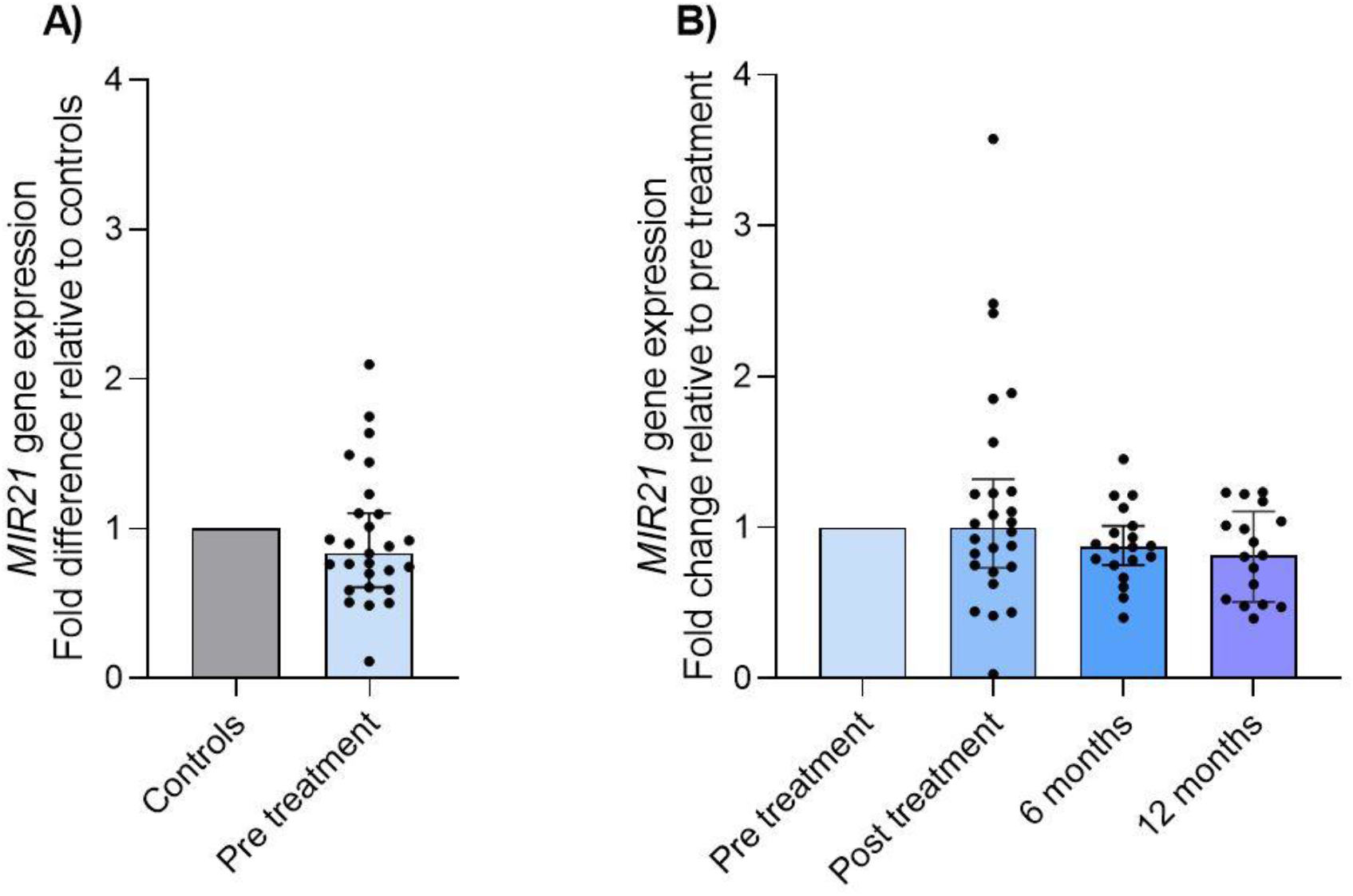
The Fold Difference or Change in Gene Expression of *MIR21* in Granulocytes from BC Patients and Healthy Controls. **A)** Fold difference in gene expression of *MIR21* in granulocytes from patients before chemotherapy normalized to the housekeeping miRNA gene *MIR24* and controls through calculation of ΔΔCt. Error bars show the interquartile range, whereas the top of the bars represents the medians. Controls are set to 1.0 as reference. Dots represent each patient (n = 27). **B)** Fold change in gene expression of *MIR21* in granulocytes from patients post chemotherapy, 6 months after chemotherapy, and 12 months after chemotherapy normalized to the housekeeping miRNA gene *MIR24* and patients pre treatment through calculation of ΔΔCt. Error bars show the interquartile range, whereas the top of the bars represents the medians. Patients pre treatment are set to 1.0 as reference. Dots represent each patient (post: n = 26, 6 months: n = 19, 12 months: n = 17).

## 4. Discussion

In this study, we show that chemotherapy induces a transient demethylation of the *VMP1*/*MIR21* region in granulocytes from postmenopausal BC patients. These changes were reversible (i.e. returned to pre-chemotherapy levels) during the first year after chemotherapy. Although we did not detect any significant associations between DNA methylation and *VMP1* or *MIR21* expression levels, demethylation of the *VMP1*/*MIR21* region may be a potential epigenetic biomarker of systemic inflammation^5,8–13^.

So far, few studies have investigated the influence of chemotherapy on DNA methylation in healthy cells and tissues from BC patients. In an epigenome-wide study, the *VMP1*/*MIR21* region was identified as the region with the largest demethylation after BC chemotherapy in PBMCs with on average 15.5 percentage points^5^. Yao et al. found the same region to be demethylated after chemotherapy in leukocytes from postmenopausal BC patients^4^. In line with these studies, we demonstrate that chemotherapy-induced demethylation of the *VMP1/MIR21* region occurs in granulocytes in addition to leukocytes^4^ and PBMCs^5^.

However, in the unique setting of the current study, we show for the first time that the chemotherapy-induced DNA methylation change is reversible. More specifically, the *VMP1/MIR21* methylation level increased on average by 9 percentage points in the BC patients 12 months after chemotherapy compared to shortly after chemotherapy, reaching a DNA methylation level that was no longer statistically different from the level before chemotherapy. This indicates that chemotherapy drives a transient demethylation of *VMP1/MIR21* in blood cells from BC patients. In the current cohort of BC patients, 83% of the patients received radiotherapy in addition to chemotherapy making it difficult to distinguish the effects of chemotherapy vs. radiotherapy. However, a previous study showed that the DNA methylation of *VMP1/MIR21* was significantly lower in chemotherapy-receiving BC patients compared to non-chemotherapy-receiving BC patients^5^, indicating that chemotherapy is the primary cause of the *VMP1/MIR21* DNA demethylation in the BC patients.

The mechanism(s) behind the demethylation of *VMP1/MIR21* in blood cells after chemotherapy treatment is not yet known. Because of the high abundance of neutrophils in granulocyte populations (>90%), the therapy-induced demethylation is most likely not explained by changes in cell type proportions. Chemotherapy is acutely immunosuppressive and negatively affects the differentiation and viability of leukocytes, however, granulocytes have a short lifespan of less than 24 hours^21^, therefore methylation differences in various neutrophil differentiation stages is most likely not the cause of the therapy-induced demethylation. Interestingly, the two oxidants H_2_O_2_ and glycine chloramine, which are released from neutrophils during inflammation, have been reported to inhibit the activity of DNA methyl transferases (DNMTs) in lymphoma cells^22^. In addition, glycine chloramine has been shown to deplete the methyl donor S-Adenosylmethionine (SAM)^22^. This indicates that the inflammatory response to chemotherapy-induced cell death can inhibit DNA methylation.

DNA methylation of *VMP1/MIR21* correlated negatively with miR-21 expression in leukocytes^14^ and PBMCs^5^ in previous studies, and an autoregulatory feedback loop has been demonstrated between miR-21 and *VMP1* expression in colorectal cancer tissues^16^. We found no significant correlation between DNA methylation and expression levels of *VMP1* or miR-21. Furthermore, neither the *VMP1* nor *MIR21* expression changed with chemotherapy in our study. The absence of correlation between *VMP1/MIR21* methylation and miR-21 expression in our study may be due to differences in study material compared to other studies. Since expression of *VMP1* and *MIR21* is low in granulocytes, an association with DNA methylation levels may not be detectable even if there is a regulatory effect.

## 5. Conclusions

With the unique follow-up samples in this study, we demonstrate that the chemotherapy-induced demethylation of the *VMP1/MIR21* region in granulocytes from postmenopausal BC patients is reversible, returning to pre-chemotherapy levels in the first year after completed chemotherapy treatment. Methylation of the *VMP1/MIR21* locus has previously been associated with inflammatory markers, and we suggest that the reduced DNA methylation after chemotherapy might be associated with therapy-induced inflammation in the BC survivors. However, further studies are needed to clarify this association and to further clarify how BC therapies impact the epigenome of healthy cells and tissues in BC survivors.

## Acknowledgements

We thank all participants who volunteered to enroll in the study. Thanks to Professor Kirsten Grønbæk at BRIC, University of Copenhagen for providing the pyrosequencing equipment. This study was financially supported by the Danish Diabetes Academy (grant number NNF17SA0031406) and the Danish Cancer Society (grant number R325-A18845)

## Abbreviations

*ACTB*: Actin beta
BC: breast cancer
miRNA: microRNA
*MIR21*: gene encoding microRNA-21
*VMP1*: vacuole membrane protein 1
PBMCs: peripheral blood mononuclear cells
PBS: phosphate buffered saline

## Notes

**Conflict of interest** All authors declare that the presented research was conducted in the absence of any conflicts of interest.

### Competing Interest Statement

The authors have declared no competing interest.

