## Supplementary data for "Chemotherapy causes a reversible decrease in *VMP1/MIR21* DNA methylation in granulocytes from breast cancer survivors"

**Supplementary table 1: Descriptive characteristics of BC patients and healthy controls.**

| | Pre treatment<br>n = 30 | Post treatment<br>n = 30 | 6 months<br>n = 21 | 12 months<br>n = 19 | Controls<br>n = 10 | p-value t-test<br>(pre vs con) |
|---|---|---|---|---|---|---|
| **Age (years)** | 59 ± 5 | 59 ± 5 | N/A | 60 ± 4 | 58 ± 4 | 0.51 |
| **Weight (kg)** | 74.8 ± 13.2 | 73.5 ± 13.0 | N/A | 74.8 ± 13.9 | 85.4 ± 25 | 0.11 |
| **BMI (kg/m$^2$)** | 26.8 ± 5.0 | 26.8 ± 4.9 | N/A | 27.2 ± 5.2 | 29.9 ± 8.8 | 0.19 |

Descriptive characteristics of healthy controls and BC patients before chemotherapy (pre treatment), shortly after chemotherapy (post treatment), 6 and 12 months after ended chemotherapy treatment. Data is presented as mean ± SD. p-values are determined from unpaired t-tests between BC patients pre treatment and controls. BMI: Body Mass Index; N/A: Not applicable.

**Supplementary table 2: Disease and Treatment Characteristics of BC Patients**

| | Number of patients (percentage) n = 30 |
|---|---|
| **Tumor stage** | |
| I | 7 (23%) |
| II | 17 (57%) |
| III | 6 (20%) |
| **Surgery type** | |
| Mastectomy | 9 (30%) |
| Lumpectomy | 21 (70%) |
| **Number of positive lymph nodes** | |
| 0 | 10 (33%) |
| 1-3 | 19 (63%) |
| 4+ | 1 (3%) |
| **ER status** | |
| Positive | 25 (83%) |
| Negative | 5 (17%) |
| **HER2 status** | |
| Positive | 13 (43%) |
| Negative | 17 (57%) |
| **Chemotherapy** | |
| Cyclophosphamide | 26 (87%) |
| Epirubicin | 23 (77%) |
| Docetaxel | 3 (10%) |
| Paclitaxel | 25 (83%) |
| Other | 1 (3%) |
| **Radiotherapy** | |
| Yes | 25 (83%) |
| No | 5 (17%) |
| **Endocrine treatment (aromatase inhibitors)** | |
| Yes | 25 (83%) |
| No | 5 (17%) |
| **Targeted therapy (Trastuzumab)** | |
| Yes | 13 (43%) |
| No | 17 (57%) |
| **Anti-resorptive treatment** | |
| Yes | 30 (100%) |
| No | 0 (0%) |

Data is presented as number of patients and percentage of the cohort in parentheses. n: sample size; ER: estrogen receptor; HER2: human epidermal growth factor receptor 2.

**Supplementary figure 1: CpG sites analyzed in the gene body of *VMP1* and promoter region of *MIR21*.**

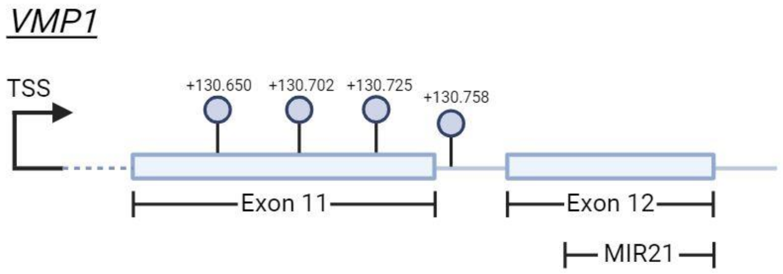


Four CpG sites in the gene body of *VMP1* were investigated through bisulfide pyrosequencing. Circles indicate the chosen CpG sites and the numbers above the circles indicate the nucleotide distance from the transcription start site (TSS). The four chosen CpG sites (+130.650, +130.702, +130.725, +130.758) are located in exon 11 and the intron between exon 11 and 12 in *VMP1* and this region has been described as the promoter region of *MIR21*, which is encoded in exon 12 of *VMP1*. Figure made in Biorender.
